# PTMProphet: Fast and Accurate Mass Modification Localization for the Trans-Proteomic Pipeline

**DOI:** 10.1101/679845

**Authors:** David D. Shteynberg, Eric W. Deutsch, David S. Campbell, Michael R. Hoopmann, Ulrike Kusebauch, Dave Lee, Luis Mendoza, Mukul K. Midha, Zhi Sun, Anthony D. Whetton, Robert L. Moritz

## Abstract

Spectral matching sequence database search engines commonly used on mass spectrometry-based proteomics experiments excel at identifying peptide sequence ions, and in addition, possible sequence ions carrying post-translational modifications (PTMs), but most do not provide confidence metrics for the exact localization of those PTMs when several possible sites are available. Localization is absolutely required for downstream molecular cell biology analysis of PTM function *in vitro* and *in vivo*. Therefore, we developed PTMProphet, a free and open-source software tool integrated into the Trans-Proteomic Pipeline, which reanalyzes identified spectra from any search engine for which pepXML output is available to provide localization confidence to enable appropriate further characterization of biologic events. Localization of any type of mass modification (e.g., phosphorylation) is supported. PTMProphet applies Bayesian mixture models to compute probabilities for each site/peptide spectrum match where a PTM has been identified. These probabilities can be combined to compute a global false localization rate at any threshold to guide downstream analysis. We describe the PTMProphet tool, its underlying algorithms and demonstrate its performance on ground-truth synthetic peptide reference datasets, one previously published small dataset, one new larger dataset, and also on a previously published phospho-enriched dataset where the correct sites of modification are unknown. Data have been deposited to ProteomeXchange with identifier PXD013210.

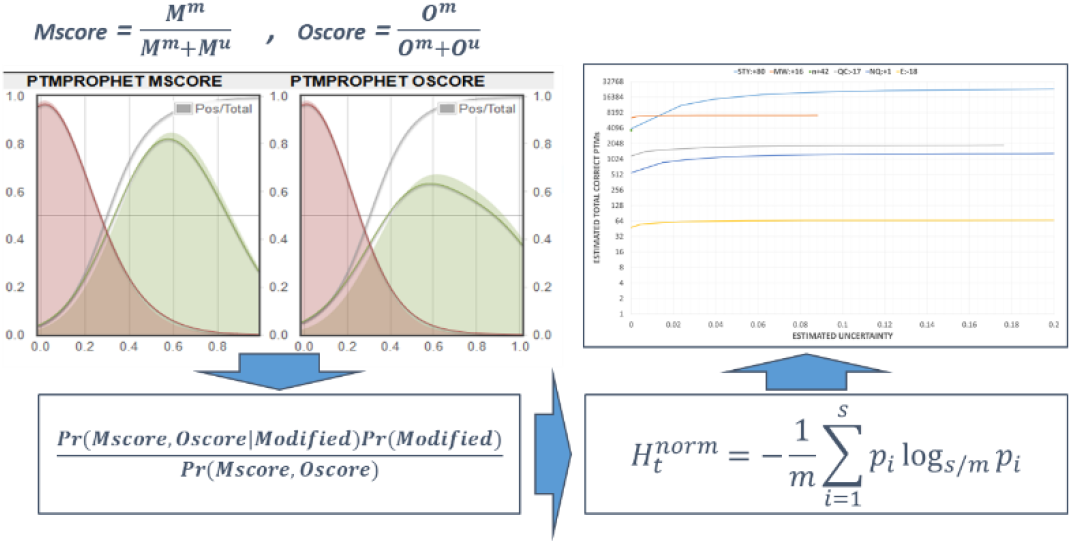

## Introduction

Proteomics utilizing mass spectrometry (MS) to interrogate the identity, quantity and modifications of proteins has become an invaluable tool to define the protein context of biological samples. Data-dependent analysis or “shotgun proteomics” remains the most widely used workflow as mature instrument workflows, software tools, and reference databases exist to identify large numbers of peptides and proteins in complex samples. This approach has given biologic insight into samples from many species and tissues. The utility of this approach extends to the detection of post-translational modifications (PTMs) and other possible modifications since all abundant ions in an experiment are analyzed together, and all modifications can be detected and characterized with suitable informatics approaches.

In a typical experiment, a large fraction of all detectable peptide ions introduced into the MS are sequentially fragmented and the resulting product ion spectra are recorded. The resultant recorded spectra from datasets that are expected to contain peptide ions are then subjected to an informatics analysis that attempts to identify the peptide sequences of the ions that yielded the spectra. However, these approaches typically require some *a priori* knowledge or assumptions on detectable proteins and potential modifications. Spectra are most commonly identified with sequence search engines such as Mascot^1^, X!Tandem^2^, Comet^3^, and MS-GF+^4^, among others. However, for each of these tools, expanding the number of potential mass modifications that must be considered expands the search space and can greatly increase computation time as well as reduce the identification sensitivity. Spectra may also be identified by spectral library searching, wherein new spectra are compared to a library of previously observed spectra. More recently, the search engine MSFragger^5^ demonstrated very fast wide mass tolerance searching that enables identification of spectra having a large mass error, potentially indicating a PTM contained within a peptide.

In each of these approaches, the correct peptide sequence is often identified and the possible location of a PTM assigned. Yet, the confidence with which the modification is localized in the event of multiple possible amino acid residues is often not well characterized. Most search engines provide the best matching peptide ion sequence and locations for PTMs, often with a confidence score quantifying the likelihood that the peptide ion sequence is correct, but do not generally provide confidence metrics for the PTM site assignments. Indeed, there are often only a few potential site determining ions that allow differentiation between a PTM located on different sites within the same peptide, and evaluating those various possibilities to derive confidence metrics is non-trivial. Robust metrics for local and global false localization rates (FLRs) should always be computed to complement the global and local false discovery rate (FDR) metrics that are currently commonly reported for peptide ion identification results. See Chalkley and Clauser^6^ for a review of FLR concepts and early tools, and Wiese *et al*.^7^ for a more recent comparison of tools.

There are several existing tools for PTM site assignment, although most of them are designed only for specific search engines or types of PTMs. Some tools determine most likely site localizations based on search engine score differences, such as the Mascot Delta Score^8^. The Protein Prospector SLIP score^9^ is built in to the Protein Prospector search engine^10^ and does not require a post-processing step. However, this score is only computed for modifications which were specified at search time; to compute SLIP scores on alternative residues not specified during the original search, a new search must be performed. Most other tools re-analyze the peaks in each spectrum to derive a PTM site localization probability based on all possible permutations of the modifications. According to the original publication, ASCORE^11^ is able to calculate localization probabilities for SEQUEST^12^ and Mascot^1^ searches of phosphopeptide data. The PTM Score algorithm re-analyzes Andromeda^13^ search results as part of the MaxQuant software package^14^. PhosCalc^15^ is an online web tool able to assess localization based on mgf or dta input files. The phosphoRS tool^16^ has been extended to handle other types of mass modifications under the ptmRS^17^ name. The P-brackets approach^18^ uses complementary product ion pairs to increase confidence in the site assignments. The LuciPHOr tool^19^ localizes phosphorylation search results from pepXML^20^ input, and LuciPHORr2^21^ extends this functionality to other kinds of PTMs. Other tools such as SLoMo^22,23^ and PhosphoScan^24^ focus on specific fragmentation types. The PTMiner^25^ tool can be used to assist in localizing the potential mass modifications discovered during open modification searches, and pSite^26^ can localize PTMs as well as compute the confidence with which de novo searching algorithms have correctly called each amino acid in a peptide sequence.

The Trans-Proteomic Pipeline^20,27,28^ (TPP) is a suite of software tools and standardized file formats that enables open, transparent and interoperable analysis of shotgun proteomics data from start to finish^29^. It is open source and designed to work on all three major operating system platforms, Microsoft Windows, GNU/Linux, and Apple MacOS X. The TPP supports many of the most popular search engine algorithms, including SEQUEST^12^, Mascot^1^, X!Tandem^2^, Comet^3^, ProteinProspector^30^, MS-GF+^4^, MSFragger^5^, among others, and enables users to apply statistical validation models that improve discrimination between correct and incorrect peptide-spectrum matches (PSMs) based on several corroborating lines of evidence that can be applied to the raw search engine scores. The iProphet tool^31^ further improves discrimination between correct and incorrect peptide identifications and enables the combination of the results of several search engines into a single result with highly controlled distinct peptide sequence level statistical significance measures. ProteinProphet^32^ applies additional modeling at the protein level to provide probabilities and FDR estimates for protein identifications inferred from the PSM and peptide evidence. In each of these tools, an expectation maximization algorithm is used to develop Bayesian mixture models from which confidence metrics are computed.

PTMProphet is a tool designed to function within the TPP software suite that models the potential sites of PTMs independently of the spectrum identification provided by a search engine, and calculates robust probabilities for each potential modification site (PMS) which provide estimates for local FLR. To elaborate with an example, if the user is searching for phosphorylations on the common amino acids of S, T and Y during search time, and decides that they also want to localize potential rare phosphorylations (e.g. on D, H and C) in phosphorylated PSMs identified by the search engine, they can evaluate these possibilities with PTMProphet without rerunning the search and by simply allowing PTMProphet to localize the additional rare amino acid modifications. In this way, the search time will not be increased by the additional complexity of the modifications considered and PTMProphet running time will be only marginally affected by the additional possibilities it will now be tasked with evaluating on a subset of PSMs.

PTMProphet also computes information content measures based on the PMS probabilities which are utilized to compute global FLR at any confidence threshold. PTMProphet reads and writes results based on the pepXML^20^ formatted output of any search engine, whether processed with PeptideProphet and iProphet or not, and results are graphically visualized in the PepXML Viewer. Below we describe the PTMProphet algorithm and demonstrate its application to several different datasets. The issue of correct assignment of PTMs underlies biologic and biomedical research where downstream research involving site directed mutagenesis and phosphomimentic amino acid substitutions is involved and time consuming. We present tools that enable a greater degree of surety in pursuing such research.

## Methods

### Implementation

As with most other major TPP analysis tools, PTMProphet is implemented in C++ that cross-compiles across Windows, Linux, and OS X into a command-line executable. This enables PTMProphet to become part of automated and distributed large-scale computing environments. The tool can also be invoked via the TPP graphical user interface (TPPGUI) for interactive use. The TPPGUI is implemented as a web interface with extensive JavaScript extensions to enhance usability. This has the advantage that the TPPGUI can easily be used locally on the same machine or on remote machines.

PTMProphet takes the results of the PSM identification process in pepXML^20^ file format as input. This will typically be the output of PeptideProphet or iProphet, but any well-formed pepXML output by any tool will work. For each PSM in the pepXML file, the original peak list is read from mzML^33^ or mzXML^34^ for subsequent analysis. The proposed peptide-ion identification for the spectrum is assessed for all user defined modification variations. Each specified type of mass modification that is expected in the peptide is assessed by considering all possible permutations of specified modifications on potentially modified sites within the peptide. For each potentially modified site on the peptide, PTMProphet compares the best scoring peptidoform with the modification occurring at that site against the best scoring peptidoform with the modification not occurring at that site and thus occurring at a different site. Peaks that are shared between two permutations provide no discrimination between the possibilities and are hence ignored. Peaks that discriminate between the different permutations are scored as described below.

The computationally intensive components of PTMProphet are multi-threaded, enabled by setting the MAXTHREADS=<number> option to a number not equal to 1 (default being 1). Multiple threads are employed primarily during the evaluation and scoring of each individual PSM, which can occur in parallel. There is no limit to the number of threads, and therefore multi-core computers can process datasets markedly faster than single core computers. Only the expectation maximization (EM) machine learning component is single-threaded since EM iterations must occur serially.

The major benefits of PTMProphet with respect to other similar tools are that it computes robust probabilities instead of arbitrary scores, computes modification information content based on the site probabilities, utilizes widely used pepXML for input and output instead of ad hoc formats, handles any combination of mass modifications not just phosphorylation, is free and open source, and is actively maintained to ensure ongoing relevance.

Although the internal format for the TPP remains pepXML, export to mzIdentML^35,36^ is also supported. Using the TPP tpp2mzid tool, the pepXML results can be exported to mzIdentML, including the PTMProphet results that are implemented using userParam XML tags specific to PTMProphet output.

PTMProphet can be downloaded as part of, and integrated in, the TPP package at http://tppms.org/. The source code is freely available at the TPP source code site https://sourceforge.net/p/sashimi/code/HEAD/tree/trunk/.

### Core Algorithm

For each PSM in the input file, PTMProphet computes the sum of all intensities of peaks that are matched given a permutation of localization as follows. Let *Ψ(P)* be the sum of all matched intensities for a given peptide *P*. For each potentially modified site *s* on *P*, PTMProphet computes:

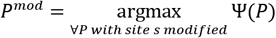

and,

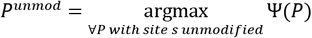

Thus, for each site *s*, *P^mod^* is the top scoring peptide, the one that maximizes *Ψ*, among all peptides that have position *s* modified. Conversely, for each site *s*, *P^unmod^* is the top scoring peptide, the one that maximizes *Ψ*, among all peptides P that have position *s* not modified.

Then, PTMProphet computes the “shared in common” matched peak intensities, termed *C(P^mod^,P^unmod^)*, for all ions identical between peptides *P^mod^* and *P^unmod^*.

Next, PTMProphet computes the discrete intensity evidence for each PSM site *s*, assuming that the site is modified, *O^m^*, and assuming the same site is unmodified, *O^u^*.

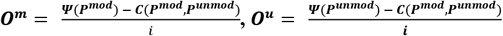

Where, *i* is the intensity of the smallest peak in the spectrum, thereby providing an automatic scaling to the smallest level of presumed noise. In order to reduce overconfidence in cases where one or two very small peaks are competing against no evidence for an alternative site, an additional, user-tunable scaling is provided by a MINO=<number> option, which adds a constant pseudo-count value (default 0) to both *O^m^* and *O^u^*. PTMProphet then computes the *Oscore* as:

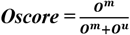

Intuitively, the *Oscore* is the fraction of the observed intensity attributable to a modification at a particular site relative to the total intensity of all site-determining peaks. PTMProphet also computes the *Mscore*, which is:

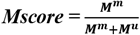

In this equation, *M^m^* is the number of peaks unique to *P^mod^*, and *M^u^* is the number of peaks unique to *P^unmod^*, i.e. not in common between *P^mod^* and *P^unmod^*. Intuitively, the *Mscore* is the fraction of peaks attributable to a modification at a particular site relative to the total number of all site-determining peaks.

PTMProphet then applies user-selectable Bayesian mixture models to all scores to compute probabilities for all sites, using a uniform prior probability for each potentially modified site. With the EM=0 option, PTMProphet estimates the probability simply using *Oscore*. With the EM=1 option, PTMProphet computes the Bayesian probability of each localization given the *Oscore*. With the EM=2 option, PTMProphet computes the Bayesian probability of each localization given the *Oscore* and *Mscore* together. With the EM=3 option, PTMProphet computes the Bayesian probability of each localization given only the *Mscore*. For options EM>0, PTMProphet applies an expectation maximization (EM) algorithm to iterate until the computed probabilities remain constant. The default setting is EM=2, which in our tests performed best, as can be seen from the comparative analysis of PTMProphet using EM modes of 0 through 3, presented in Supplemental Table 3

Next, all probabilities are renormalized so that for each PSM the sum of the probabilities for each type of mass modification equals the total number of modifications. For example, if a particular peptide with five possible modification sites contains two phosphorylated residues, the sum of all five probabilities will equal 2.0. The modifications are potentially moved from the original location placed by the search engine to the locations with the highest probabilities, which can occur when the search engine is unable to correctly localize the modification at search time due to spurious peaks or other algorithmic deficiencies to discern between closely related modified peptides. Peptide sequence strings with all embedded probabilities are also written out to pepXML in a format encoding a probability to three decimal places following each residue. For example, in a PSM with one phosphorylation and one oxidation site, a string such as S(0.000)EM(0.780)M(0.220)EEDLQGAS(1.000)QVK can be written in the pepXML, denoting the probability, *p_i_*, that each potential site, *i*, harbors a particular type of modification. For this example, the probabilities for phosphorylation are *{ 0.000, 1.000}* and for oxidation are *{0.780, 0.220}* at the corresponding potential sites of those modifications.

### Comparing Modified PSMs

Site probabilities are only directly comparable between PSMs having the same number of modifications and the same number of sites. Yet in a standard experiment it is natural to expect a diverse population of peptides to be present having different numbers of potentially modified sites and different numbers of modifications. Therefore, a problem arises when we consider comparing modified PSMs identified within a single dataset between themselves and between different datasets. Intuitively, a site probability of 0.75 in a peptide with one phosphorylated residue and two potentially phosphorylated sites (e.g., S(0.250)ENNEEDLQGAS(0.750)QVK), suggests that there is a strong possibility (three quarter chance) that the modification is present at the site with the higher probability, which is greater than random chance (50%) that the modification is present at that site. Alternatively, if there are three modifications and four sites, e.g., S(0.750)ES(0.750)S(0.750)EEDLQGAS(0.750)QVK, the same probability 0.75 means we have no additional information regarding the localization, since in this PSM a site probability value of 0.75 is exactly the random chance expectation given the number of modifications and potential sites present in the peptide.

To address this issue, the normalized per-modification information content of the PTM site assignment probabilities, *I_t_*, can be used instead of site probabilities to directly compare PSMs, regardless of the number of modifications and modification sites they contain. Computing this for all PSMs provides a value by which all PSMs can be directly compared, sorted and filtered. Based on the *I_t_* values, estimates of the uncertainty contained within a filtered modified PSM list can be measured. Computing these values for different datasets and analysis methods provides a method by which these datasets and methods can be evaluated and directly compared in the interest of optimizing methodologies. Computing the information content of modified PSM site assignments is described in the following section.

### Information Content Calculation

As a final step, PTMProphet computes the Normalized Per-modification Information Content, *INFO_t or I_t_*, of each PSM and for each type *t* of modification present in the PSM, as follows:

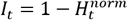

Where, 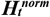 is the normalized per modification entropy^37^ of modification of type *t*, having the range *[0, 1]*. It is calculated according to the formula:

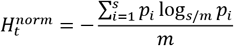

Where:

***t*** is the modification type
***s*** is the number of potential sites of modification of type ***t*** in the peptide
***m*** is the number of modifications of type ***t*** in the peptide
***p_i_*** is the PTMProphet probability of the modification occurring at site ***i***

Additionally, the number of modifications of type *t* that can be localized with certainty in a given PSM, *LMODS_t* or *M_t_*, has the range *[0, m]* and is calculated by PTMProphet as:

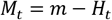

Where:

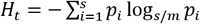

For example, a value of 2.94 might indicate that nearly 3 modifications of type t are localized with high certainty in a given PSM.

In addition to information content, PTMProphet computes and reports the mean best PTM-site probability for each PSM and modification type it contains. For each type of modification localized in each PSM, this feature calculates the mean PTM-site probabilities of the top *m* number of highest confidence sites within the PSM, of that modification type. As we demonstrate below, the mean best probability statistic for modifications of type t, *MBPr_t*, provides a remarkably accurate estimate of PSM-level false localization rates (FLR).

### PepXML Viewer

The TPP PepXML Viewer application has been enhanced to display PTMProphet results graphically, allowing for visual exploration of PTMProphet localization data. Figure 1 provides an example of a series of PSMs from the published reference phosphopeptide dataset (described below) for the peptide THLGTGMERSPGAMER. There are two types of modifications considered in this analysis, phosphorylation (S,T,Y) and oxidation (M,W.) Columns 1-3 in Figure 1 show the iProphet probability, the PeptideProphet probability, and the peptide sequence including the graphical display of the PTMProphet results. Columns 4-6 report the computed mean best probability, normalized information content, localized modification quantity for the localization of modified S,T,Y residues for each peptide. Columns 7-9 show the same for the localization of modified M,W residues. The final column in Figure 1 is the spectrum identifier; additional columns showing other peptide properties such as count of enzymatically consistent termini, retention time, precursor intensity among others can be displayed in the web interface but are not shown here.

**Figure 1.**
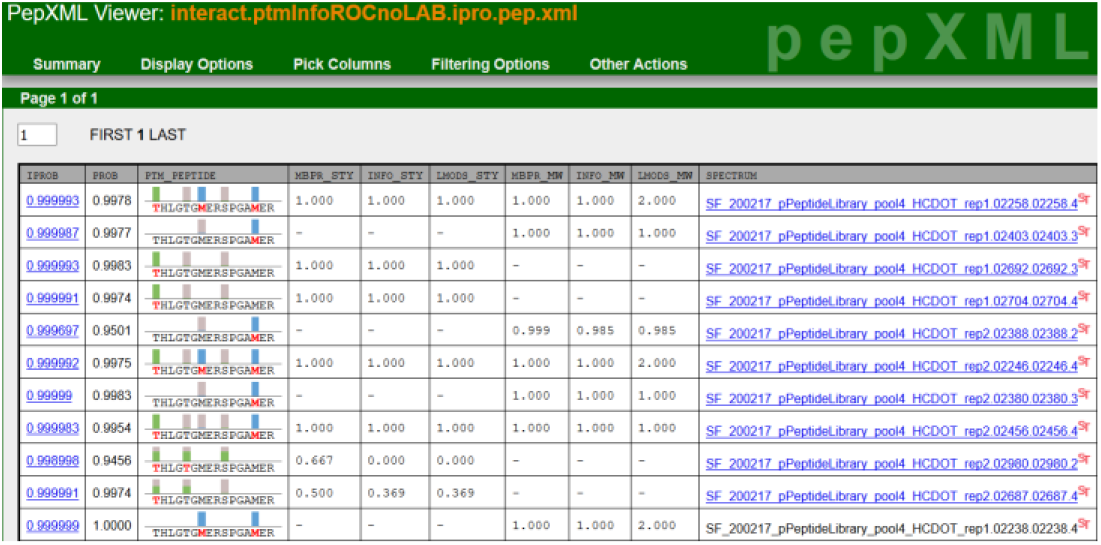
PepXML Viewer graphical display of PTMProphet results from the previously published reference dataset for peptide THLGTGMERSPGAMER. In the third column of this display, bars are placed above each residue that is a potential site for a mass modification. Gray indicates probable absence at a site, while a non-gray color indicates probable presence at that site, proportional to the height of the box. A full height box indicates a probability of 1. Hovering the mouse pointer over a bar reveals a tooltip with extended numerical information. Each type of modification has a different color. See text for further discussion of the figure.

In the graphical display of column 3 (Figure 1), each potential site has a bar above the residue. A gray bar indicates zero probability at a potential site, while the height of a non-gray colored bar indicates the fractional probability of the mass modification being correctly localized to that site. Each type of mass modification has a different color (in this figure, green is phosphorylation and blue is oxidation, although this is not fixed.) The amino acid residue letter where the mass modification is assigned (due to the highest probability) is colored red. The example peptide THLGTGMERSPGAMER has three potential phosphorylation sites and two potential oxidation sites. In the first row (Fig.1), the phosphorylation is confidently localized to the N-terminal threonine, the first amino acid residue, and both methionines are oxidized. The second PSM (row 2) has no phosphorylation at all and the last methionine only is oxidized. The PSMs in rows 3 and 4 have high-confidence phosphorylation localization on the first residue and no oxidation. The PSM in row 9 has minimal peak evidence to localize two present phosphorylated residues, and the probability at all three sites is 0.667; notice that the modification localization information content of this peptide is zero. For the PSM in row 10, a single phosphorylation is present on either the first or second threonine with probability 0.500, but presence on the serine is ruled out and given a probability of 0.000.

## Results

To demonstrate the performance of PTMProphet, we ran the tool against three example datasets, i) a small reference dataset of synthetic peptides previously published and used to evaluate confident site localization with an Thermo Fisher Scientific Orbitrap Fusion Tribrid mass spectrometer (Dataset #1, Ferries *et al*. 2017)^17^, ii) a large in-house generated and unpublished dataset of synthetic phosphopeptides with known phosphorylated sites (Dataset #2), and iii) a phospho-enriched cell lysate dataset known to contain many mass modifications (Dataset #3, Söderholm *et al*.)^43^.

For dataset #1, we compare the performance of PTMProphet to the originally published results, which we further analyzed for information content of the modified site assignments. First we assessed the accuracy of the PTMProphet implementation with this dataset consisting of a synthetic phosphopeptide library containing 175 distinct peptide sequences with 191 phosphorylation sites. The synthesized peptides are divided into five pools for data dependent LC-MS/MS analysis on a Thermo Fisher Scientific Orbitrap Fusion Tribrid mass spectrometer, with the isomeric phosphopeptides (where the same peptide sequence is phosphorylated at different positions) allocated to different pools. The original data were generated with several different fragmentation regimes (HCD, EThcD, and neutral-loss-triggered ET(ca/hc)D) and mass analyzers for MS/MS (Orbitrap (OT) versus ion trap (IT)). Data acquired by all of these collection methods are supported by PTMProphet. For simplicity, we only discuss the HCD results with high mass accuracy MS1 and MS2 scans acquired in the Orbitrap, which was reported by the authors to be the best performing method in terms of numbers and rates of correctly localized phosphosites identified by the ptmRS analysis.

The RAW files for Dataset #1 were obtained from the ProteomeXchange^38,39^ repository PRIDE^40,41^ identifier PXD007058 and converted to mzML^33^ with the ProteoWizard^42^ msConvert tool with peak picking enabled. The high mass accuracy MS2 spectra were searched both with Comet and X!Tandem with the high-resolution K-score^43^ (HRK) enabled. Mass tolerances were set to 25 ppm for precursors and 0.02 *m*/*z* for fragment peaks, along with an allowance for semi-tryptic peptides and up to 4 missed cleavages (important because some of the synthetic peptides have more than 2 missed cleavages.) In general, it is not very common to see 4 missed cleavage peptides and most natural datasets are only searched with 2 missed cleavages, unless the sample is only lightly digested. It is only important in this particular dataset because some of the synthetic peptides have this many cleavages. This results in a slight increase in search time, but does not affect the running time of PTMProphet. The searches were performed allowing for fixed modification of carbamidomethyl on C, and variable modifications for M, W oxidation and S, T, Y phosphorylation. The Comet and X!Tandem search results were individually processed with PeptideProphet using high mass accuracy setting in ppm mode, semi-parametric modeling, and an option to not penalize deltaCn* results for Comet (indicating homology between top matching peptides to a spectrum, a common occurrence for PSMs with mass modifications, where the top matches are the same peptide sequence with different modified sites.) The two resulting files were merged and modeled with iProphet to produce a single pepXML output file. The resulting single pepXML file was processed through PTMProphet with default settings, except for MAXTHREADS=0 (allowing processing of spectra to occur in parallel on all cores in the machine), and specified mass modifications STY=79.9663 and WM=15.9949.

For each PSM, mass offset and tolerance parameters are learned automatically by PTMProphet from the data by applying a Bayesian mixture model algorithm for mass tolerance selection. Output was written to a new pepXML file with updated localizations along with probabilities and models. Based on the learned iProphet models, applying an iProphet probability threshold of 0.9 yields a global peptide level FDR of 0.004, correctly identifying 4125 PSMs (17 incorrect), with a sensitivity of 0.90. Decoy counting shows there are 13 decoy PSMs passing this threshold, matching 8 unique peptide sequences, out of 4142 PSMs mapping to 1225 distinct peptidoforms and 609 distinct peptide sequences. Of these PSMs, 1467 mapped to a peptide precursor matching one or more peptidoforms in the peptide mixture, both in sequence and number of modifications. Of these PSMs, we excluded those that could have been a result of potential carryover from a prior run, which left 1420 PSMs, which were then used to compute FLR statistics as follows. The PSMs were sorted based on phosphorylation localization mean best probability (MBPr) in descending order. The resulting list was traversed in order of decreasing MBPr, computing the global average error and the global reference-based FLR among all results, including the current PSM and those above it in the list. We also computed the number of correctly localized PSMs and the associated FLR rates per the reference. Figure 2 panels A and B summarize the results obtained from this analysis. In Figure 2A, we can see the number of correctly localized PSMs at different FLRs as measured by counting the PSMs with correctly and incorrectly localized sites according to the reference. In Figure 2B, we can see the comparison of PTMProphet estimated FLRs to the reference-based FLRs at the PSM level. In general, we can observe from Figure 2B that the PTMProphet analysis of this dataset gives a conservative FLR estimation for the entire dataset, meaning the mean best probability measure can be used to estimate false localization errors that are slightly above the actual error based on the reference annotation of correctly and incorrectly localized PSMs. Furthermore, based on this analysis and as displayed in Figure 2A, PTMProphet correctly localized 1164 PSMs at a reference-based PSM FLR of less than 1% in this dataset.

**Figure 2.**
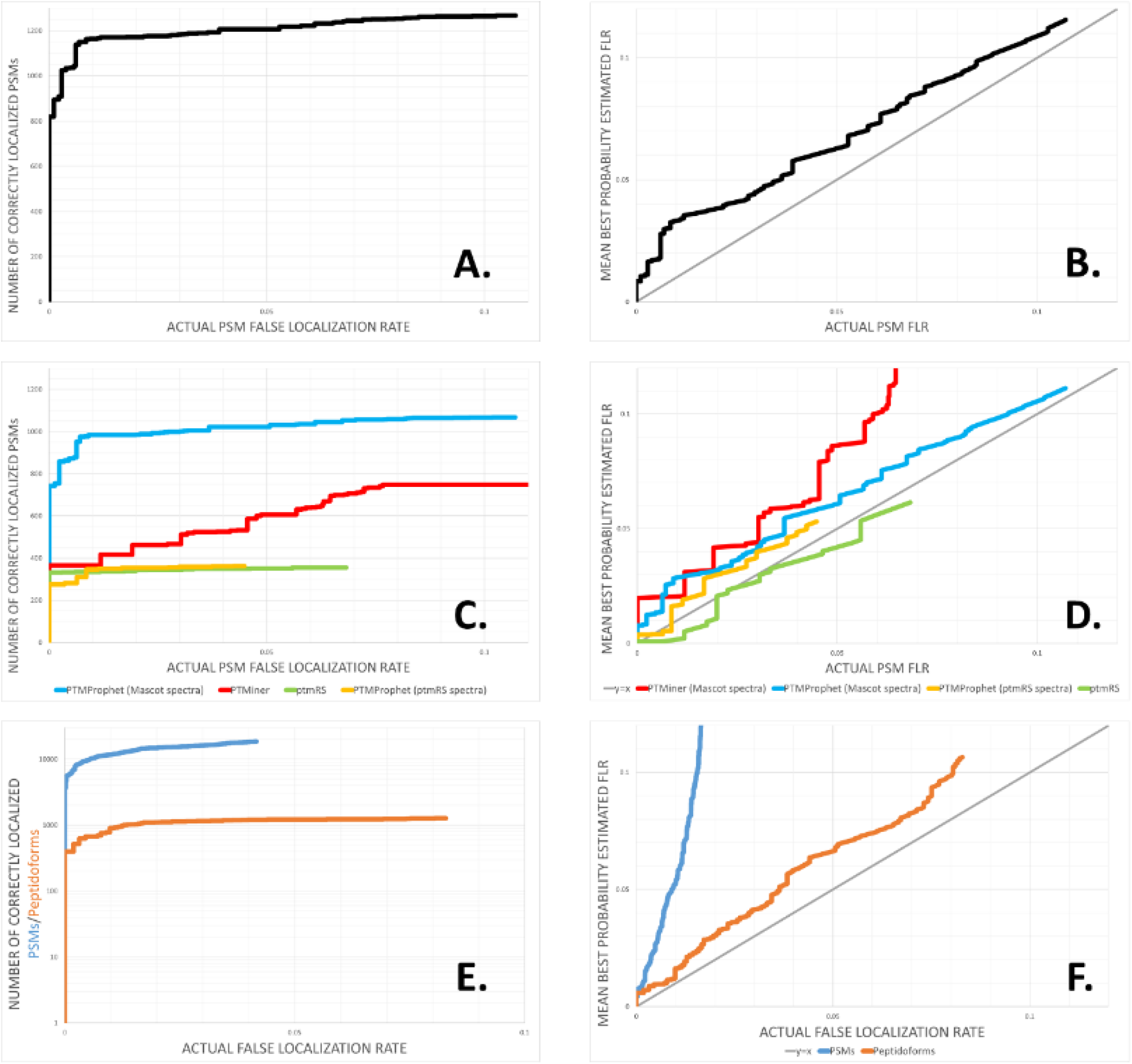
Panels A and B are from the Comet+X!Tandem TPP-based analysis of Dataset 1. Panel A displays a ROC plot showing the reference-based PSM FLR on the x-axis plotted against the number of correctly localized PSM on the y-axis; Panel B displays a plot comparing the reference-based PSM FLR on the x-axis vs the mean best probability estimated PSM FLR on the y-axis, the grey diagonal indicates the line where the reference-based FLR equals the estimated FLR. Panels C and D show the comparison of the Mascot analysis localization results of ptmRS in green, the PTMProphet localization analysis on all Mascot PSMs in blue, the PTMProphet localization analysis only on those Mascot PSMs for which ptmRS returned a localization result in yellow, and PTMiner results in red. Panel C displays ROC plots showing reference-based PSM FLR on the x-axis plotted against the number of correctly localized PSMs on the y-axis; Panel D displays the plot comparing the actual reference-based PSM FLR on the x-axis to the mean best probability estimated PSM FLR on the y-axis, the grey diagonal indicates the line where the reference-based FLR equals the estimated FLR. Panels E and F display the analysis results from running PTMProphet analysis on Dataset 2. Panel E displays ROC plots showing reference-based FLR on the level of PSMs in blue and best scoring peptidoform in each run in orange on the x-axis plotted against the number of correctly localized PSMs and best scoring peptidoforms in each run on the y-axis; Panel F displays the plot comparing the actual reference-based PSM and best scoring peptidoform in each run FLR on the x-axis to the mean best probability estimated PSM and best scoring peptidoform in each run FLR on the y-axis, the grey diagonal indicates the line where the reference-based FLR equals the estimated FLR.

Next, we compared the performance of PTMProphet to the originally published results of this dataset. While we can’t compare PTMProphet in detail to all available site localization tools with their different strengths and weaknesses, we compare it here to ptmRS^17^ and PTMiner, a pair of representative recently developed and well-performing PTM localization tools. We wanted to compare PTMProphet to the best performing tools and thus used the best performing dataset from the best performing method, HCD Orbitrap, and analysis tool, ptmRS, from the Ferries *et al*. 2017 publication. The analysis includes filtered Mascot search results containing 3563 PSMs and filtered ptmRS results containing 542 PSMs. We ran PTMProphet on the Mascot results using the same settings that we applied to the Comet/X!Tandem/TPP analysis. Because ptmRS results are reported as potential modification site probabilities, for each PSM, we were able to easily apply the formulas for computing *MBPr_STY_* and *I_STY_* to ptmRS phosphorylation site assignments. We computed these statistics for each localized PSM, applying it to PTMProphet, ptmRS, and PTMiner analysis of the published Mascot results.

The performance pattern observed for the PTMProphet analysis of the Mascot result is similar to the performance pattern observed for PTMProphet on the Comet + X!Tandem analysis of the same dataset. Figure 2D demonstrates that PTMProphet and PTMiner computed localization errors yield conservative FLR estimates when compared with reference-based error rates. In comparison, the ptmRS computed localization errors yield FLR estimates that are anticonservative when compared with reference-based error rates, meaning that ptmRS is underestimating the true error rate. The ptmRS tool gives an anticonservative result for nearly all reference-based FLRs in the dataset. For instance, at the reference-based FLR of 1%, PTMProphet estimated FLR is slightly above 1%, while ptmRS estimated FLR is still below 0.25%. Figure 2C demonstrates that on a small subset of PSMs ptmRS does identify a greater number of correctly localized PSMs at reference FLRs of less than 1%. At FLRs above this PTMProphet shows a slightly higher number of correctly localized PSMs than ptmRS. In all, when we consider the number of correctly localized peptidoforms at an actual FLR of less than or equal to 1% we find that PTMProphet identified 156 peptidoforms, ptmRS identified 148 peptidoforms and PTMiner identified 141 peptidoforms in this data set (data presented in Supplemental Table 1.)

The analysis described above was applied only to those PSMs which matched the order sheet sample peptides for each run, both in sequence and number of modifications. In other words, we only analyzed PSMs where the correct precursor was in the correct sample, excluding PSMs with incorrect number of modifications or those that had potential for cross-run carryover contamination. Cross-run contaminating peptides could be correctly localized, but present in the incorrect run and thus represent false negative results that should be left out from the ground-truth answer assessment. Among the PSMs where the correct precursor was found in the correct sample and the PSM could not have been the result of cross-run carryover, the ptmRS site assignment disagreed with the reference on 26 PSMs (2 of which were high confidence), of which 16 were correctly assigned by PTMProphet (13 with high confidence and 3 with mid-range confidence.) PTMProphet results considering only the PSMs for which ptmRS also had a result and filtered in a similar fashion, disagreed with the reference on 17 PSMs (3 of which were high confidence), of which 7 were correctly assigned by ptmRS (all 7 with high confidence). Thus, PTMProphet showed less disagreement with the synthetic reference peptides and was able to correctly localize more PSMs among those where the ptmRS software disagreed with the reference set. Supplemental Table 1 lists the PSMs that were incorrectly localized by each software.

Supplemental Table 1 provides the worksheets that were used to generate the figures for PTMProphet and ptmRS analyses of the localization results of the Ferries *et al*. dataset. The table includes Excel exported raw results of PTMProphet and ptmRS that were processed to generate the plots shown above. Also included is the comparative analysis of the incorrectly localized PSMs by ptmRS, PTMiner, and PTMProphet, as compared to the other software tools.

Next, we evaluated PTMProphet on a larger dataset (Dataset #2 has been deposited to PRIDE^44^ under ProteomeXchange^39^ identifier PXD013210) generated from 1342 chemically synthesized phosphopeptides with 5329 S, T and Y residues that could potentially be phosphorylated (see Supplemental Table 2). These peptides are up to 26 amino acids long and contain between one and eleven S, T and Y residues per sequence, with 83% of all peptides containing two to six S,T,Y sites. This reference set contains 493 distinct peptide sequences (37%) and 849 distinct peptidoforms (63%) with the same sequence but phosphorylated residues assigned to different sites on the sequence, e.g., SIS[167]TVSSGSR and SISTVS[167]SGSR, to reflect the challenges of correct site assignment in natural samples. 629 of the 1342 selected human peptides have empirical evidence while 713 peptides were designed *in silico* by taking backbones from empirical peptides and moving the empirically observed phospho group to another S, T or Y residue. Peptides were pooled as such some isomeric peptides were allocated in different pools while others were in the same pool, again to reflect the challenges in real samples. This dataset was collected from pools of 95 peptides (except for 4 pools with fewer peptides) by data dependent LC-MS/MS analysis on a SCIEX TripleTOF 5600+ (see Supplemental Methods for details).

The raw data were converted to mzML using the SCIEX mzML converter with option /proteinpilot for peak picking, /zlib for compression and /index for indexing the files. The data were searched using both Mascot (version 2.4) and X!Tandem (version Jackhammer TPP 2013.06.15.1). The searches were done specifying parent mass tolerance of 0.1 Da with isotopic offsets for both search engines and fragment mass tolerance of 0.8 Da for Mascot. Carbamidomethyl on Cys was the only static modification applied in the search and the variable modifications include phosphorylation on S, T and Y and oxidation on M for both search engines. The spectra were searched against a database of these synthetic peptides appended to a large synthetic peptide database described in Kusebauch *et al*. ^45^ and decoys based on the randomized versions of the synthetic peptides.

The search results of both search engines were processed with PeptideProphet and the pepXML results of both PeptideProphet analyses were combined using iProphet (compiled from SVN revision 7336 of the TPP code). Finally, we ran PTMProphet on this dataset to localize the PTMs with default settings, except for MAXTHREADS=0 (allowing processing of spectra to occur in parallel on all CPU cores), MINPROB=0.9 (making PTMProphet apply localization to PSMs having an iProphet probability of 90% and higher.) Next, we analyzed the FLRs comparing PTMProphet estimated FLRs based on mean best probabilities to the actual FLRs based on correctly localized PSMs and peptidoforms.

As can be surmised from Figure 2 panels E and F, when comparing estimated FLRs to actual FLRs, PTMProphet provided a conservative estimate of false localization error on both the level of PSMs and on the level of peptidoforms, taking the highest scoring PSM identifying each peptidoform type in each run. The highly conservative nature of PSM level FLR on this dataset stems from the fact that there is a large number of PSMs in the dataset when compared to the number of peptidoforms in the dataset. PTMProphet assigns the sites correctly on many of the lower quality PSMs but with lower probability than in the higher quality PSMs, so while the localization is correct the lower PTM site probability on the lower scoring PSMs contribute to an upper bound estimate on the actual localization error. As can be observed when comparing the orange curve to the blue curve in Figure 2F, when we considered only the best scoring PSM of each peptidoform in each run, the FLR estimates became much more accurate when compared to the dataset ground truth, while still remaining conservative estimates of site localization error.

We also applied PTMProphet to Dataset #3, a phosphopeptide enriched dataset derived from human cells. We selected and reanalyzed the PRIDE repository identifier PXD001620, as originally presented and described in detail by Söderholm *et al.^46^*. Briefly, the dataset is from human macrophage cells infected with influenza A virus. The protein samples are digested with trypsin, enriched for phosphopeptides using PHOS-Select™ Iron Affinity Gel (Sigma Aldrich, MO, USA) and subjected to data dependent LC-MS/MS analysis on a Q Exactive mass spectrometer. The raw files were converted to mzML using msConvert and searched with Comet and X!Tandem, followed by PeptideProphet analysis on the search engine results. The search mass tolerances were set to 10 ppm for precursors and 0.03 m/z for fragment peaks, along with an allowance for semi-tryptic peptides and up to 2 missed cleavages. The database used in the search was the PeptideAtlas Tiered Human Integrated Search Proteome^47^ (THISP) database compiled on 2018-01-01, appended with the Influenza A protein sequence database. The THISP protein sequences were of Tier 2, meaning the sequence database includes the core primary isoforms from neXtProt (nP), cRAP contaminants, plus “varsplic” alternative splice isoforms from neXtProt, immunoglobulin variable region sequences from Swiss-Prot, and IMGT^48^. Also appended were decoy sequences created from random shuffling of tryptic peptides in the target protein sequences. The searches were performed allowing for fixed modification of carbamidomethyl on cysteines and variable modifications as follows: methionine and tryptophan oxidation, serine, threonine and tyrosine phosphorylation, n-terminal acetylation, pyro-glu from glutamine and pyro-carbamidomethyl as a delta from carbamidomethyl-cysteine, pyro-glu from glutamic acid, and deamidation of asparagine and glutamine. To keep running times for searches at a reasonable level, the selection of modifications considered in the search was based on the modifications used in the original publication, with the additions of deamidation of asparagine and glutamine, and the oxidation of tryptophan, which our laboratory regularly uses in database searching. The Comet and X!Tandem search results were individually processed with PeptideProphet using high mass accuracy setting in ppm mode, semi-parametric modeling and enabling the option to use the expectation score for modeling of Comet results. The two resulting PeptideProphet pepXML files were merged and modeled with iProphet to produce a single pepXML output file.

The resulting single pepXML file was processed through PTMProphet with default settings, except for MAXTHREADS=0 (allowing processing of spectra to occur in parallel on all CPU cores), MINPROB=0.8 (making PTMProphet apply localization to PSMs having an iProphet probability of 80% and higher, corresponding to an iProphet estimated PSM error rate of 0.4% and PSM sensitivity of 97.9% at this iProphet probability threshold) and considering the following mass modifications: STY=79.9663, WM=15.9949, QC=-17.026549, E=-18.010565, NQ=0.984016, and n-terminus=42.0106. The selection of MINPROB is largely an arbitrary choice made by the user. This should be made based on the corresponding error-rate the user is willing to accept in the results. We selected a lower minimum probability on this dataset where the correct localizations are unknown, using some lower scoring PSMs to extend the FLR range in the ROC plots, while keeping the PSM error rate well below 1%. Figure 3 summarizes the modifications localized with varying degrees of certainty, across all PSM matches in the dataset, based on the information content calculations for each type of modification analyzed by PTMProphet. We can see from the plot, matching expectation based on the experimental design, the most common modification observed in this dataset is indeed phosphorylation. PTMProphet was able to localize in total, across all PSMs: 16,876 phosphorylation modifications of S, T or Y, 7220 oxidation modifications of M or W, 3717 n-terminal acetylation modifications, 1784 pyroglu from Q or pyro-carbamidomethyl from carbamidomethyl-C modifications, 1156 deamidations of Q or N modifications, and 65 pyro-glu from E modifications, at 5% estimated FLR or less. The n-terminal acetylation modification is always localized with perfect certainty for the obvious reason that each linear peptide has but a single n-terminal location.

**Figure 3.**
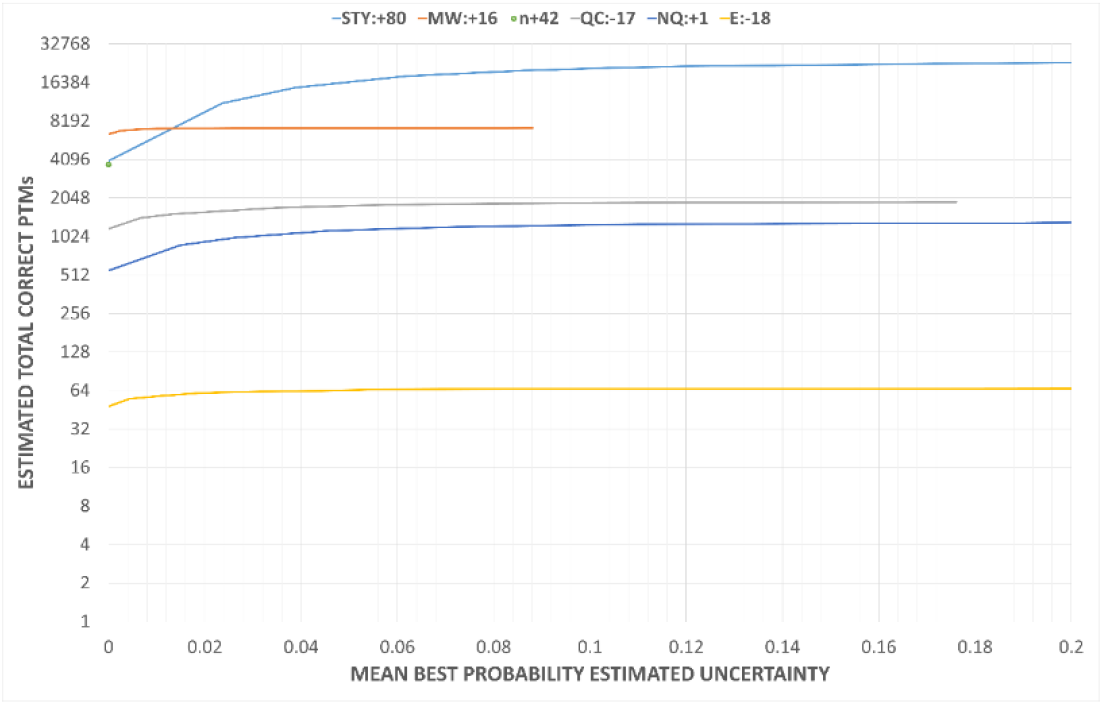
Estimated total number of correctly localized modifications of each type versus the mean best probability (MBPr) estimated false localization rate (FLR), computed by summing (1-MBPr) over all PSMs traversed in order of decreasing MBPr.

## Conclusion

In molecular cellular biology and biomedicine the correct and appropriate assignment of a post-translational modification enables the rapid progression of further investigations on molecular pathology and physiology. The investigator needs surety on mass spectrometry observations to progress work via routes such as phosphomimetic site directed mutagenesis. We therefore developed the peptide modification localization confidence tool PTMProphet (with supporting tutorials) in an automatable fashion through its inclusion in the Trans-Proteomic Pipeline for the rapid and accurate calculation of localization statistics for mass spectrometry PSM data. Use of information content for PTM site assignments is incredibly useful for sorting, filtering and comparing of modified PSMs. PSMs containing different numbers of potential PTM sites and different numbers of PTMs can now be directly and fairly compared. Additionally, the information content provides an upper-bound FLR estimate that can be used to directly compare site assignments made by different parameters or algorithms as long as the algorithm reports its localization as site probabilities that sum to the number of modifications contained in the peptide. Although, in our analysis the use of information content for PTMProphet site assignments provided an ultra-conservative upper-bound estimate of actual FLR and mean best probability provided a less conservative though still conservative result, it appears the actual statistic to apply depends on the algorithm used and less conservative algorithms may get more accurate results by employing information content calculations. Whichever way FLR is ultimately estimated, we think that it would be beneficial for PTM site assignment algorithms to report their localization confidences as PTM site probabilities that sum to the number of PTMs in each PSM, as opposed to arbitrary scores.

PTMProphet can be applied to any modification that is measured as a mass difference in a mass spectrometer. For modifications that are generally labile in a collision cell, e.g. glycosylation, special considerations must be made. Specifically, collision energy and selection of fragmentation methods must be carefully considered when attempting to measure labile PTMs. For PTMs that appear on a specific amino acid (e.g. lysine in the case of ubiquitination), special considerations for enzyme selection during sample prep must be made to avoid trypsin cleavage of ubiquitinated lysine residues. PTMProphet localizes modifications of any type, as specified by the user during execution time.

In order to facilitate adoption, we provide a step-by-step tutorial of the tool that guides users how to use and evaluate the software on a sample dataset on their own computer. The tutorial is available at http://www.tppms.org/tools/ptm/, and leads the reader through installing TPP on a Microsoft Windows computer, downloading the sample data set, processing the sample data set, and evaluating the results. As can be understood from running the tutorial, improved ease of use of the software, from the installation of the TPP in general, to downloading of public data, to the invocation of PTMProphet, and to visualizing, filtering and sorting of PTMProphet site assignments specifically, has been achieved. In the current version of the TPP, the PTMProphet software tool is available as open source code that is compiled and provided freely within the TPP architecture. Following the installation of the TPP, the user has to copy their own data to a specified location readable by the TPP on the analysis computer, select this data inside the application, and specify a minimal set of required parameters, to be able to run the analysis. Selection of user specified parameters are minimized to specifying the types of modifications to be localized in the data. For each PSM, PTMProphet will automatically select the mass differences to be applied based on the machine learned mass difference parameters for that PSM. Thereby, the burden to run the software and select optimized parameters for analysis is minimized for the user.

PTMProphet software scales well so it can be applied to many datasets and has been demonstrated to work well on previously published and new complex datasets with comparison to other well-characterized PTM localization software. Its performance and usability capabilities on very large datasets is demonstrated by the identification and localization of all 8 million PSMs represented in the Human Phospho PeptideAtlas 2017 being processed through a development version of PTMProphet to derive global phosphorylation patterns across dozens of datasets. All future versions of the Phospho-PeptideAtlas will utilize this version or updates of PTMProphet to enhance the quality of the large-scale phosphopeptide data identification repository of PeptideAtlas.

## Supporting information

Supporting Information Summary

## Acknowledgements

This work was funded in part by the National Institutes of Health, National Institute of General Medical Sciences grants: R01GM087221, R24GM127667, P50GM076547, the National Institute of Allergy and Infectious Diseases grant: R21AI133335, the National Institute of Biomedical Imaging and Bioengineering grant U54EB020406, and the National Heart Lung and Blood Institute grant: R01HL133135, the National Institute On Aging of the National Institutes of Health under Award Number U19AG023122, Medical Research Council (ADW), and by the Cancer Research UK major centre award (20761). We would also like to thank Jimmy Eng, Andrew Keller and Alexey Nesvizhskii for the meaningful discussions that helped bring this project to fruition.

## Supporting Information

Supplemental Figure 1 - Analysis Workflow

Supplemental Figure 2 – PTMProphet Usage

Supplemental Methods: Methods for Generating Dataset #2.

Supplemental Table 1: Comparison of ptmRS Analysis to PTMProphet Analysis of Dataset #1.

Supplemental Table 2: List of Synthetic Phosphorylated Peptides in Dataset #2.

Supplemental Table 3: PTMProphet comparative analysis of EM options 0, 1, 2, and 3

## Notes

The authors declare no competing financial interest.

