## Supporting Information Summary for "PTMProphet: Fast and Accurate Mass Modification Localization for the Trans-Proteomic Pipeline"

| **File** | **Description** |
| --- | --- |
| SupplementalFigures.pdf | Supplemental Figure 1 - Analysis Workflow  Supplemental Figure 2 – PTMProphet Usage |
| SupplementalTable1.xlsx | PTMProphet, ptmRS and PTMiner analyses for Dataset #1 |
| SupplementalTable2.xlsx | PTMProphet analysis of identified PSMs and proteoforms from Dataset #2 |
| SupplementalTable3.xlsx | PTMProphet comparative analysis of EM options 0, 1, 2, and 3 |
| SupplementalMethods.docx | Methods by which Dataset #2 data was generated |
