## Supplemental Figures for "PTMProphet: Fast and Accurate Mass Modification Localization for the Trans-Proteomic Pipeline"

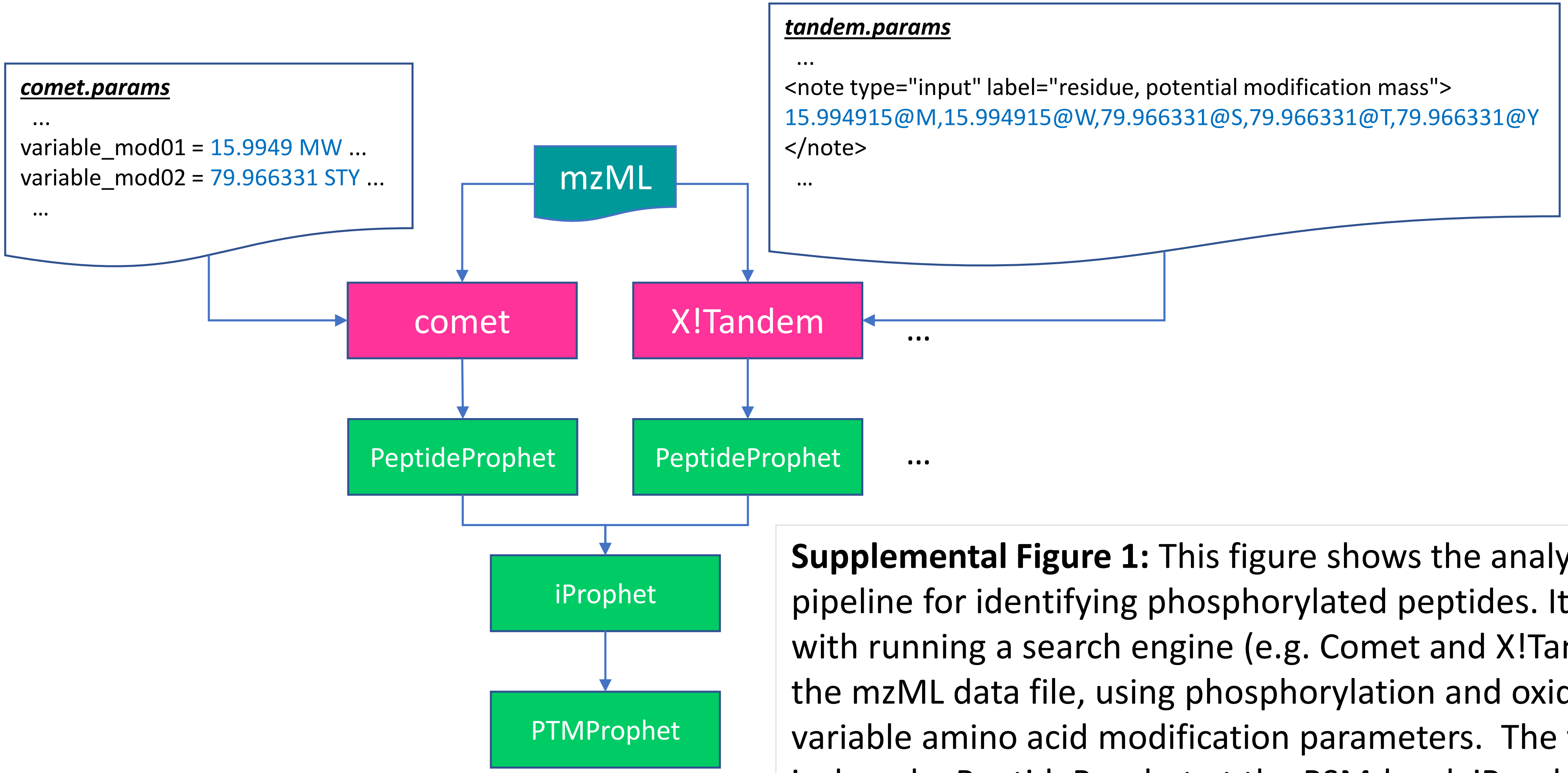


**Supplemental Figure 1:** This figure shows the analysis pipeline for identifying phosphorylated peptides. It starts with running a search engine (e.g. Comet and X!Tandem) on the mzML data file, using phosphorylation and oxidation as variable amino acid modification parameters. The validation is done by PeptideProphet at the PSM-level, iProphet at the peptide-level and PTMProphet at the PTM-lever.

***PTMProphetParser Usage:***

**USAGE**: PTMProphetParser <OPTIONS> <input_file.pep.xml> [<output_file>]

**OPTIONS**

**NOUPDATE** Don't update modification_info tags in pepXML

**EM**=<number>: Set EM models to <number> which can be 0, 1, 2, or 3 (*default: 2*) :
0 -> no EM,
1 -> Intensity EM Model Applied Only,
2 -> Intensity and Matched Peaks EM Models Applied,
3 -> Matched Peaks EM Model Applied Only

**KEEPOLD** Option to retain old PTMProphet results in the pepXML file (*default: off*).

**VERBOSE** Option to produce WARNINGS to help troubleshoot potential PTM shuffling or mass difference issues (*default: off*).

**STATIC** Use static fragppmtol for all PSMs instead of dynamically estimates offsets and tolerances (*default: off*).

**FRAGPPMTOL**=<number> When computing PSM-specific mass_offset and mass_tolerance, use specified default +/- MS2 mz tolerance on fragment ions (*default: 15*).

**PPMTOL**=<number> Use specified +/- MS1 ppm tolerance on peptides which may have a slight offset depending on search parameters (e.g. iTRAQ4plex) (*default=1*).

**MINPROB**=<number> Use specified minimum probability to evaluate peptides (*default=0.9*).

**MAXTHREADS**=<number> Use specified number of threads for processing (*default=1*).

**MASSDIFFMODE** Treat the mass difference between measured and theoretical mass as a modification and localize

**LABILITY** Compute Lability of PTMs

**DIRECT** Use only direct evidence for evaluating PTM site probabilities

**IFRAGS** Use internal fragments for localization (*default: do not use internal fragments*)

**AUTODIRECT** Use direct evidence when the lability is high, use in combination with LABILITY

**MAXFRAGZ**=<integer> Limit maximum fragment charge (*default: precursor charge, negative values subtract from precursor charge*).

**MINO**=<integer> Use specified number of pseudo-counts when computing Oscore (*default=0*).

**NOMINOFACTOR** Disable MINO factor correction when MINO= is set greater than 0 (*default: apply MINO factor correction*)

**NIONS**= Use specified N-term ions, separate multiple ions by commas (*default: a,b for CID, c for ETD*)

**CIONS**= Use specified C-term ions, separate multiple ions by commas (*default: y for CID, z for ETD*)

**MASSOFFSET**=<number> Adjust the massdiff by offset <number> (*default: 0*)

**<amino acids, n, or c>:<mass_shift>:<neut_loss1>:...:<neut_lossN>,<amino acids, n, or c>:<mass_shift>:<neut_loss1>:...:<neut_lossN>**
Specify modifications (*default: STY:79.9663,MW:15.9949* )

**Supplemental Figure 2:** This figure shows the complete USAGE statement of the PTMProphet software and default parameters, in yellow, utilized by PTMProphet.
