## Supplemental Methods for "PTMProphet: Fast and Accurate Mass Modification Localization for the Trans-Proteomic Pipeline"

**Synthetic phosphopeptides (example data set 2)**

1342 phosphorylated peptides were individually chemically synthesized as free amine at the N-terminus and carboxylic acid at the C-terminus (JPT Peptide Technologies, Micro-scale Peptides, 50 nmol per peptide, crude). Phosphorylated serine, threonine and tyrosine were introduced as phosphorylated building block and cysteine residues as carboxyamidomethylated cysteine building block.

Peptides were analyzed in pools of maximum 95 peptides on a SCIEX TripleTOF® 5600+ equipped with a Nanospray-III® Source (Sciex) and an Ekspert™ nanoLC 425 with cHiPLC® system operated in trap-elute mode (Eksigent). Peptides were loaded on a cHiPLC trap (200 µm x 500 µm ChromXP C18-CL, 3 µm, 120 Å) and washed for 10 minutes at 2 µL/min. Peptides were eluted on a nano cHiPLC column (75 µm x 15 cm ChromXP C18-CL, 3 µm, 120 Å) with 0.1% formic acid in water (A), 0.1% formic acid in acetonitrile (B) (v/v) at 300 nL/min using a gradient from 3% to 43% B in 80 min, 43%-63% B at 80-85 min and 63%-83% B at 85-87 min at a flow rate of 300 nL/min. A survey scan (TOF-MS) was acquired in the *m/z* range of 400-1,250 Da with 250 msec accumulation time. The 30 most intense precursor ions with charge state 2-5 were selected for fragmentation with rolling collision energy. MS/MS fragment spectra were collected in the range of 100-1,500 Da with 50 msec accumulation and a 1.8 s period cycle time. Former target ions were excluded for 15 seconds.
